# Redefining the structure of tip-links in hair-cells

**DOI:** 10.1101/2022.08.19.504503

**Authors:** Veerpal Kaur, Sanat K. Ghosh, Tripta Bhatia, Sabyasachi Rakshit

**Affiliations:** Department of Chemical Sciences, Indian Institute of Science Education and Research Mohali, Punjab, India; Department of Physical Sciences, Indian Institute of Science Education and Research Mohali, Punjab, India

**Author notes:** Correspondence to Sabyasachi Rakshit.

## Abstract

Tip-link, the gating-spring in hearing, modulates the mechanoelectrotransduction process and protects the hearing sensory machineries from overexcited auditory inputs. Tip-links are formed by two exceptionally long cadherins, cadherin-23 and protocadherin-15. It is proposed that both the cadherins first independently form cis-dimers through lateral interactions and finally, engage in trans-interactions to form the heterotetrameric assembly in tip-links. In fact, the molecular details of the protocadherin cis-dimers are well-understood. However, little structural and molecular details are known for independent cis-dimer of cadherin-23. Using photo-induced cross-linking of unmodified proteins in solution and on lipid-membranes, we observed no trace of cadherin-23 cis-dimers, rather we captured trans-homodimers of cadherin-23. We thus, redefine the tip-link structure where protocadherin-15 remains as cis-dimers, however, two individual cadherin-23 binds to protocadherin-15 cis-dimer for tip linking.

## Introduction

Tip-links, a crucial element of the mechanoelectrical transduction (MET) process in hearing, are present in the inner ear of humans and other vertebrates^1,2^. Hair cells in the inner ear possess bundles of stereocilia atop, arranged in a staircase manner. Each stereocilium in the bundle is connected to a neighboring shorter stereocilium at the tips through tip-links. The base of the tip-link on shorter stereocilium is connected to a mechanoelectrotransducer (MET) channel^3,4^. In the hearing, auditory inputs deflect stereocilia which subsequently, generate mechanical tension at tip-links. Tip-links in response conveys a threshold force to the MET channel for opening and thus, convert the mechanical input into an electrical signal^5,6^. Tip-links in this process of mechanotransduction serve as low-force pass-filter as gating-spring which transduce a small force from auditory inputs and filter overexcitation to protect the sensory machineries^7^.

Tip-links are made up of two long non-classical cadherins, cadherin-23 (Cdh23) and protocadherin-15 (Pcdh15)^8,9^. Cdh23 is among the longer protein having 27 extracellular domains (EC domains), contributing 2/3 to the length of the tip link^8^. Pcdh15 has 11 EC domains followed by membrane-proximal PICA domain^10,11^. From the electron micrographs of stereocilia, the tip-links appeared as helical threads with a length of 120-170 nm and thickness of 5 nm^12^. The lower part of the helical coil is formed by Pcdh15. In situ imaging of stereocilia using cryo-electron microscopy and antibody-modified Au-nanoparticles staining clearly depicted the intertwined lateral dimers of Pcdh15^13^. The residues at the EC3 domain and the membrane-proximal PICA domain that drive the lateral interactions in Pcdh15 were also identified from X-ray crystallography of varying EC-domains ^10,14^. Finally, a series of mutations including Leu138Asp, Leu138Ala,, Arg198Ala, Asp255Lys, Val250Asp,, Lys345Ala, and Lys345Asp were made to pin-point the residues through the disruption of the lateral interactions^14^. Capturing the independent lateral dimer of Cdh23, however, has been inconsistent^8^. Rather, the homophilic trans-dimer of Cdh23 is well established^15^, and their physiological role in cell-cell adhesion is also well-understood^16–18^. Thus, questions arise on the existence of the lateral dimers of Cdh23, either in solution or on the membrane of stereocilia. Here, we employed the Photo-Induced Cross-Linking of Unmodified Proteins (PICUP)^19^ in a native form to capture the lateral dimers of Cdh23, even if the interactions are weak. We testified the efficiency of our method with the lateral dimer of Pcdh15 and the transdimer of Cdh23. From our work, we re-define the structure of tip links.

## Results

### Cdh23 EC 1-27 exists as a monomer whereas Pcdh15 EC 1-11 PICA as a dimer in solution

Cdh23 is a large protein of molecular weight ∼ 325 kDa so its expected dimer is at ∼ 650 kDa. Traditional methods such as size exclusion chromatography and native gel electrophoresis are not a convenient way to detect the ∼ 650 kDa dimer of Cdh23. To date, most of the studies are based on electron micrography to visualize the dimer of such large proteins but in these chances of artifacts are more as samples are fixed and stained for imaging. Other ways to probe proteinprotein interactions are chemical cross-linking but these chemical methods suffer from poor reaction yield and the requirement of large-scale modification in side chains of proteins that can lead to artifacts in results. To detect the dimer of Cdh23 and Pcdh15 in the solution we used the photo-induced cross-linking of unmodified proteins (PICUP) method^19^ **(Figure 1A)** which is a time-saver, does not require protein modifications, and can capture short-lived stable dimers or oligomers formed in solution as well as on membrane. Cdh23 and Pcdh15 both belong to the cadherin superfamily, a large calcium-dependent family of proteins. It’s also known that calcium concentration is critical for proper mechanotransduction in hair cells, so we performed PICUP experiments at varied calcium concentrations. The recombinant protein constructs that we used for the PICUP are Cdh23 with entire EC domains (Cdh23 EC 1-27), Pcdh15 with all EC domains and PICA domain (Pcdh15 EC 1-11 PICA), and Pcdh15 with two outermost EC domains (Pcdh15 EC1-2), with GFP-6xHis at the C-terminus of all.

**Figure 1.**
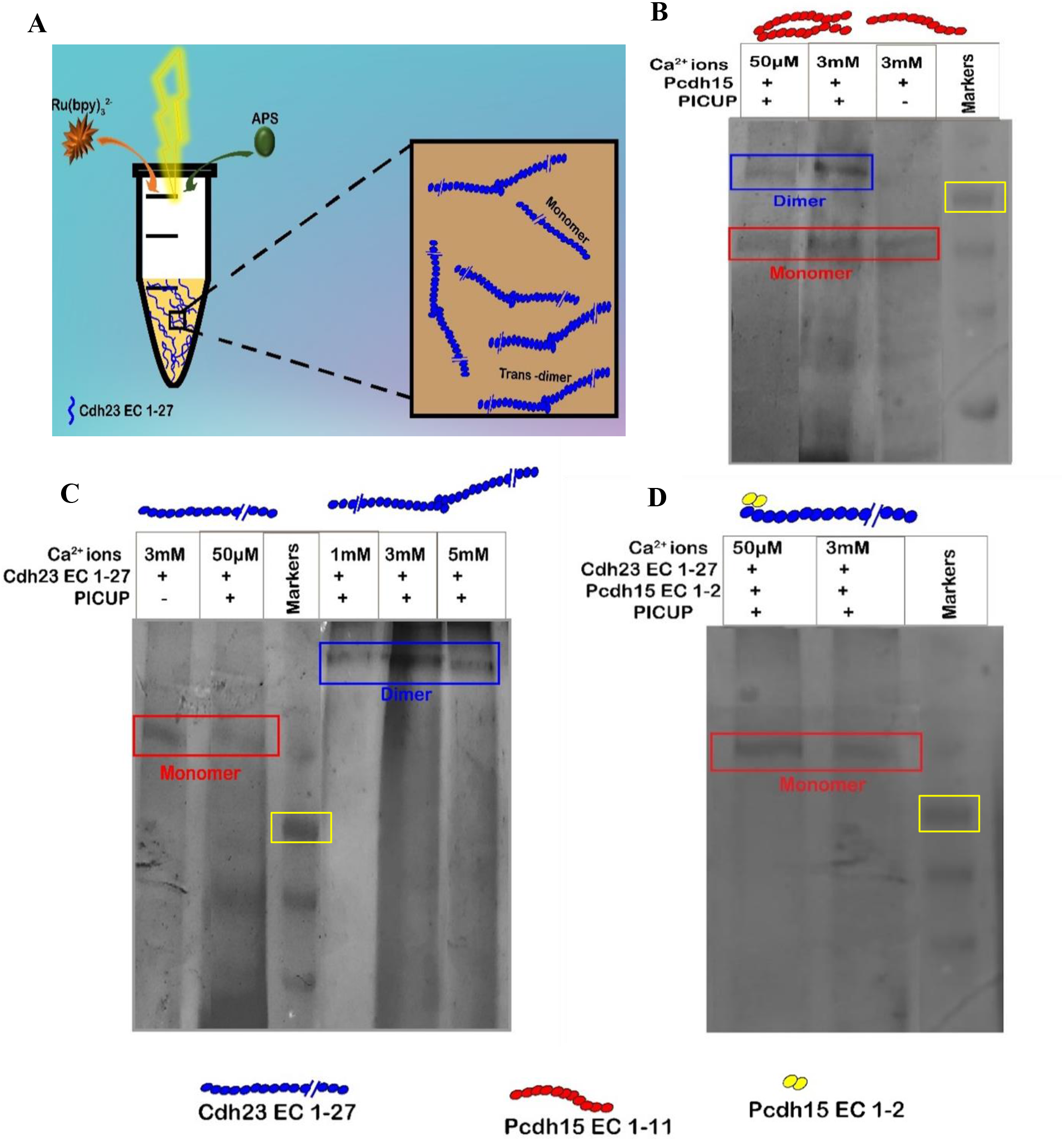
Photoinduced coupling of unmodified proteins in solution. **(A)** Schematics of PICUP protocol. **(B)** Silver stained SDS PAGE of Pcdh15 EC1-11 PICA in solution after PICUP at two extreme [Ca^2+^] of 50 µM and 3 mM. For comparison, a no PICUP protein is run at extreme right before the ladder. The red-colored bead structures at the header of the PAGE represent Pcdh15 EC1-11 PICA in different complex association. **(C)** Silver stained SDS PAGE of Cdh23 EC 1-27 in solution after PICUP at 1mM, 3mM, and 5mM of [Ca^2+^], respectively. The -blue colored bead structures at the header of the PAGE represent Cdh23 EC1-27 in different complex association. **(D)** Silver stained SDS PAGE of Cdh23 EC 1-27 after PICUP, where the trans-interaction site is pre-blocked by Pcdh15 EC1-2 (yellow beads). In B, C, & D, ‘No PICUP’ condition is indicated as (-) and with PICUP as (+). The blue rectangle marks the corresponding dimer, red rectangle marks the monomer, and the yellow marks the ladder band at 315 kD, respectively.

Cis-dimer of Pcdh15 is now well-characterized. The dissociation constant (*K*_*D*_) of the homophilic cis-dimer of Pcdh15 in solution^14^ is 1.8 µM. As proof of the PICUP protocol, and subsequent detection method, we first performed PICUP on 3 µM (above *K*_*D*_) of Pcdh15 EC 1-11 PICA at endolymph [Ca^2+^] (50 µM) and high-[Ca^2+^] of 3 mM, respectively. Subsequently, we ran SDS-PAGE followed by silver staining. As expected, we observed homophilic cis-dimer bands of Pcdh15 at ∼340 kDa in both the calcium concentrations **(Figure 1B)**. Interesting to note that no dimer band was visibly detected in no PICUP condition. This is expected as often denaturation breaks structure-specific interactions. Further, the protein bands in low-[Ca^2+^] were faint than the 3 mM [Ca^2+^] and this could be due to differences in staining for different conformations of the protein. Cadherins are known to adopt [Ca^2+^]-dependent conformations.

We next performed PICUP of Cdh23 EC1-27 at varying [Ca^2+^] of 50 µM, 1mM, 3mM, and 5 mM. We used an optimal concentration of 6 µM for Cdh23 EC1-27 for the PICUP as beyond this we always observed smearing of bands in the SDS-PAGE. Notably, at higher concentrations (> 4 µM), Cdh23 EC1-27 undergoes liquid-liquid phase separation^20^. We observed dimer bands at ∼650 kDa **(Figure 1C)** in all calcium concentrations under PICUP, except at the endolymph [Ca^2+^] of 50 µM. It is important to note that the dimer band at ∼650 kDa does not confirm the complex orientation, cis-dimer, or trans-dimer. Cdh23 forms trans-homodimer with a dissociation constant^15^ of ∼18 µM and mediates cell-cell adhesion^18^. In order to confirm the configuration of the dimer band of Cdh23, we planned to repeat the PICUP after blocking the trans-interactions. Cdh23 forms the heterophilic trans-complexes with Pcdh15 EC1-2 with higher affinity than the homophilic trans-complexes. We, therefore, re-performed the PICUP of Cdh23 EC 1-27 but in presence of Pcdh15 EC 1-2 (5 µM) in the buffer **(Figure 1D)**. Interestingly, we did not observe any trace of dimer for Cdh23 EC 1-27 even at two extremes of [Ca^2+^] of 50 µM and 3 mM, respectively. Since the detection limit of the silver staining is ∼2 to 5 ng/protein band in polyacrylamide gel, it is, thus, implied that no fraction of Cdh23 EC1-27 exists as cis-dimers in solution.

### No cis-dimers of Cdh23 on artificial lipid membrane

A lipid membrane limits the diffusion of anchored molecules in two dimensions, and thus, influences the binding affinity of proteins through cooperative interactions^21^ and membrane fluctuations^22,23^. Cadherins are trans-membrane proteins and are naturally confined for lateral diffusions. Moreover, Cdh23 engages in weak transient interactions and forms cis-clusters prior to even any trans-mediated localization at the cell-cell junction^20^. We, thus, hypothesized that the cis-clustering of Cdh23 on the membrane may assist the lateral dimerization of Cdh23. Accordingly, we prepared (Giant Unilamellar Vesicles) GUVs using a mixture of dioleoylphosphatidylcholin (DOPC) and 1,2-dioleoyl-sn-glycero-3-[(N-(5-amino-1-carboxypentyl)iminodiacetic acid)succinyl] (nickel salt) (DGS NTA(Ni^2+^))^24^, and attached cadherins (Pcdh15 and Cdh23, respectively) through C-terminal 6xHis tags **(See Methods)**. The attachments of the individual cadherins were confirmed from the C-terminal GFP tags **(Figure 2A)**. Since free diffusion of proteins tethered to the membrane is essential for lateral proximity, and so lateral complexation, we confirm the diffusion of proteins from FRAP experiments on GUVs tethered with Pcdh15 and Cdh23, respectively^20^ **(Figure S1, Table S1)**.

**Figure 2.**
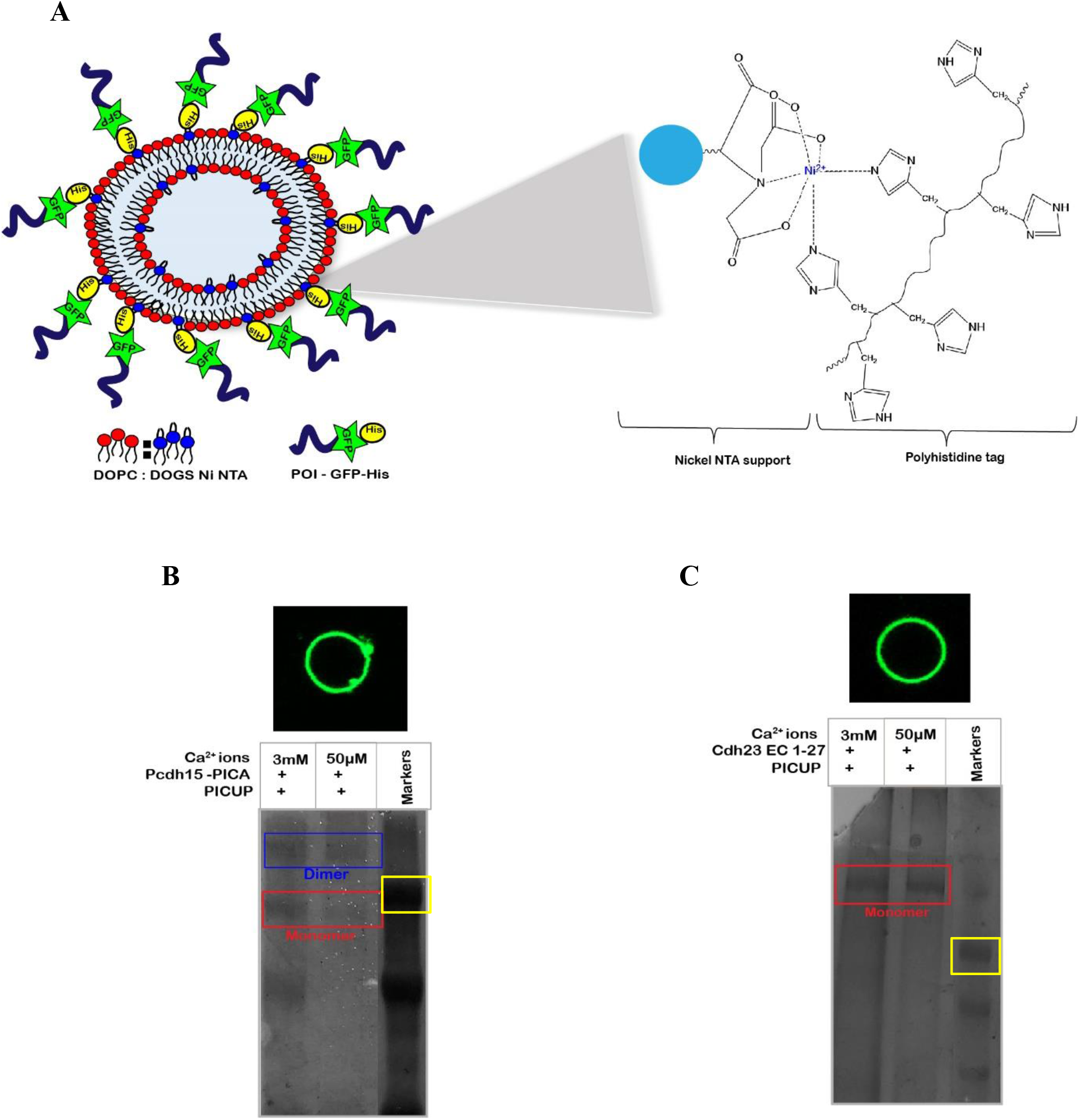
PICUP on membrane-anchored proteins. **(A)** Conceptual representation of membrane-anchored proteins, tethered to the membrane through Ni-NTA vs 6xHis-protein interactions. **(B)** Fluorescence image of a GUV homogenously tethering Pcdh15 EC1-11 PICA. The green color is from the GFP tagged Pcdh15. The bottom panel describes the Silver stained SDS-PAGE of Pcdh15 on the membrane after PICUP at two [Ca^2+^]. **(C)** Silver-stained SDS PAGE of Cdh23 EC 1-27 after PICUP on a membrane at 50µM and 3mM [Ca^2+^], respectively.

To trap the stable oligomers in the protein cluster on the membrane, we performed PICUP on GUVs. For detection of oligomeric distributions, the GUVs after PICUP were incubated with imidazole to elute the membrane-anchored proteins into solution, followed by centrifugation and gel-electrophoresis. Here again, we first used Pcdh15 as proof of the protocol, and identified the co-existence of lateral-dimer and monomer of Pcdh15 EC1-11 PICA even in the clusters, at 50µM and 5mM of [Ca^2+^], respectively **(Figure 2B)**. However, we did not observe any trace of lateral dimer for Cdh23 EC1-27 **(Figure 2C)**, indicating that Cdh23 EC1-27 exists as a monomer even on a membrane. The weak transient interactions that mediate cis-clustering of Cdh23 on the membrane, cannot influence stable cis-dimerization.

## Conclusion

The *in-situ* electron micrograph of stereocilia clearly depicted the thread-like arrangement of tip-links^12^,^25^. Subsequently, myriads of studies have been employed to identify the conformational and structural origin of the thread-like tip-links ^26^. Soon it was revealed that the constituent cadherins, Pcdh15 and Cdh23, may exist as lateral dimers and form the helical-shaped tip-links. In support, the thread-like assembly for Pcdh15 cis-dimer is also identified ^14^, and subsequently, the molecular interactions are deciphered ^10^. On the same note, the molecular detail of the Cdh23 lateral-dimer is still under exploration. Meanwhile, electron micrographs of Cdh23 depicted a large heterogeneity in the dimeric assembly. Structural elucidations of small EC domains of Cdh23 also proposed the probable lateral-interaction sites on Cdh23. However, no single studies report the stable and predominant existence of Cdh23 cis-dimer. On the contrary, the trans-dimer of Cdh23 is established long ago along with their molecular details. Overall, a continuous delay in capturing and deciphering the lateral-dimer of Cdh23 raised serious concern about its existence.

Using a photo-induced cross-linking method of natively folded proteins, we established that Cdh23 does not have any detectable affinity towards lateral dimerization, unlike Pcdh15. The tetrameric assembly of tip-link is thus predominantly guided by the cis-dimer of Pcdh15. We propose that the evolution of such a tetramer is kinetically-driven. Tip-links are dynamic in nature and thus, undergo rapid binding-unbinding cycles. While the tetrameric assembly delays the unbinding process, binding of a dimer protein with another dimer protein in a sterically confined region can retard the process. We speculate that the existence of Cdh23 as a monomer on a membrane thus facilitates the binding of tip-links during regeneration **(Figure 3)**.

**Figure 3.**
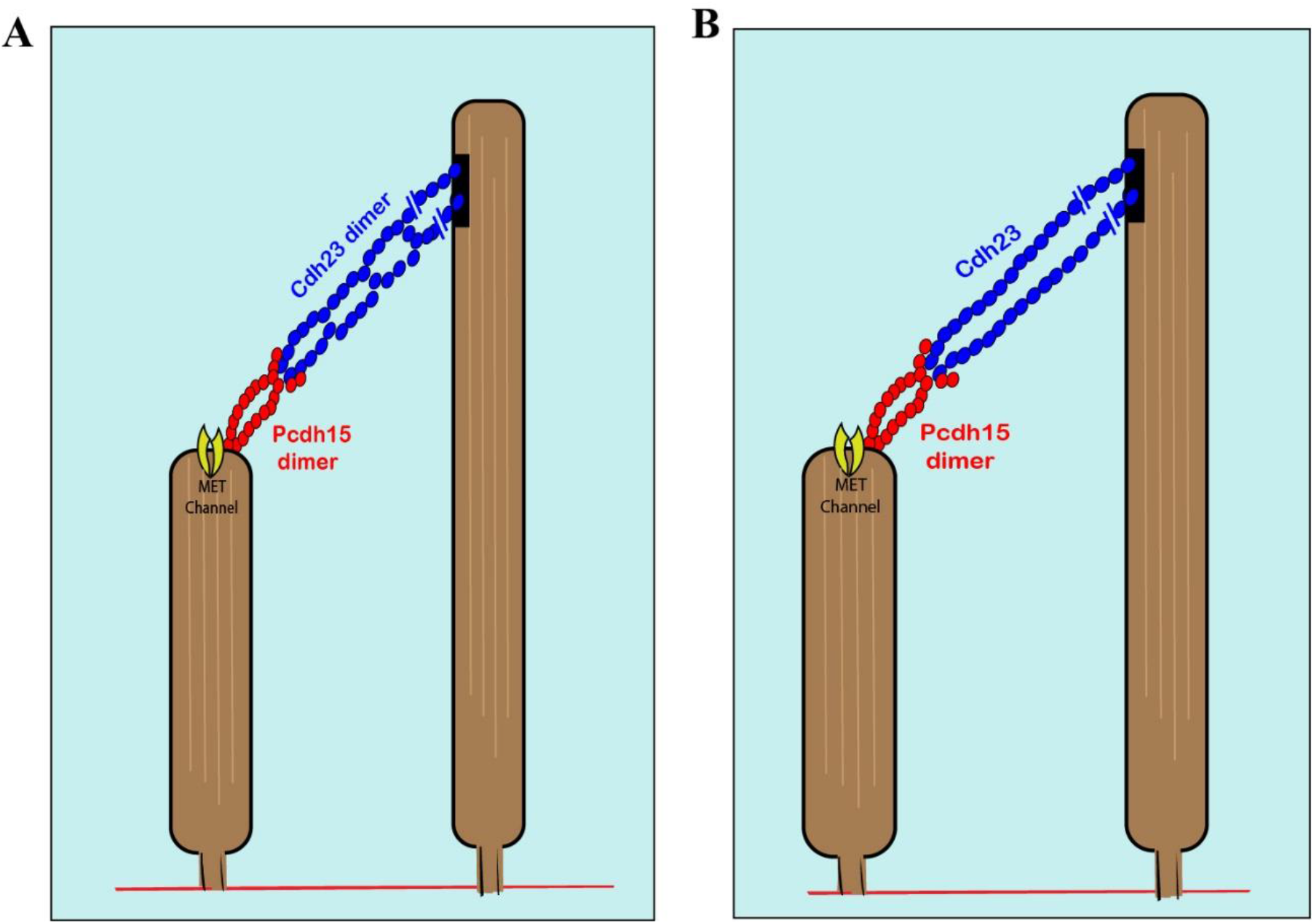
The correction in the tip-link structure. Scheme **(A)** Depicts the existing model of tip-link. Scheme **(B)** Represents the redefined tip-link structure indicating no lateral dimerization of Cdh23 in the tip-link.

## Material and Methods

### Protein expression and purification

Plasmids of Cdh23 EC 1-27, Pcdh15 EC 1-11 PICA, and Pcdh15 EC 1-2 with 6x His and GFP tag at the C-terminus were transfected and expressed in the ExpiCHO suspension cell system (A29129 ThermoFisher Scientific) using the standard prescribed protocol. After 7 days, cells were harvested and media was pellet down at 2000 rpm for 15 min at room temperature. Subsequently, the media was collected and dialyzed in the dialysis buffer (25 mM HEPES, 150 mM NaCl, 100 mM KCl, and 2mM CaCl_2,_ pH 7.5) for about 48 hours. The dialyzed media was then purified with Ni -NTA-based affinity chromatography to elute the purified proteins. The purity and presence of protein were confirmed using 6% SDS-PAGE and western blotting with specific antibodies against GFP, Cadherin-23, Pcdh15, and His-tag. Desired calcium concentrations were maintained by dialysis of the purified protein with a dialysis buffer of varying calcium concentrations.

### PICUP in solution

During photoinduced coupling, reaction volume was maintained up to 20 µl having 18 µl of 6 μM purified Cdh23 EC 1-27 or 18µl of 3μM Pcdh15 EC 1-11 with 1 µl of electron donor solution (150 mM NaCl, 15 mM sodium phosphate, 0.125 mM [Ru(bpy)_3_Cl_2,_ pH=7.5]) and 1 µl of an electron acceptor, 2.5 mM of Ammonium persulfate (APS). This 20 µl reaction volume in a transparent vial was irradiated by visible light for 0.5 sec followed by quenching of the radical reaction with ∼7 μl of SDS-gel loading dye (4x). SDS-PAGE and silver staining were done to detect the PICUP results. Pcdh15 EC1-2 (5 μM), the blocker of trans-homodimerization of Cdh23 was added during PICUP experiments.

### GUVs preparation

GUVs with lipid composition 97.5 % DOPC (Sigma-Aldrich): 2.5 % DGS NTA Ni (Sigma-Aldrich) in chloroform was prepared using gel-assisted GUV formation method^24^. 5% (w/w) solution of PVA (sigma) was prepared in the water while continuous stirring and heating at 90°C. 200-300 µl PVA solution was coated on a microscope coverslip followed by drying at 50 °C in the oven for 30 min. PVA-coated coverslips were washed with MQ water to remove extra PVA and dried for 30 min at 50 °C. 20-30 µl of lipids were spread on a dried PVA-coated surface and placed in a vacuum for 1 hour to evaporate chloroform. Finally, 500-600 µl sucrose (350 mM) was added to a coated coverslip for 1 hour to harvest GUVs. Harvested GUVs were collected into an eppendorf.

### Protein attachment to GUVs and PICUP of membrane-tethered proteins

GUVs formation was confirmed by labelling with Nile red dye. After that GUVs were incubated with desired protein in appropriate concentration for 7-8 hours at 4°C. Protein attachment on GUVs was confirmed from GFP signals.18 µl of protein tethered GUVs were subjected to the PICUP condition mentioned above. To elute out the proteins from the membrane, PICUP treated samples were incubated with 250mM imidazole for 10 min. The result was observed on SDS-PAGE followed by silver staining.

### FRAP and data analysis

Fluorescence recovery after photobleaching was monitored at the polar region of tethered GUVs at 63X magnification using a super-resolution microscope (ZEISS LSM 980 Airyscan 2).

We measured the normalized fluorescence intensity in the selected photobleached region for three independent sets of experiments and plotted it against the time. After global fitting of the fluorescence intensity data with recovery time using the equation, f (t) = A (1 – e ^-τ t^), where A indicates the mobile fraction of proteins on the membrane and from the slope, we find τ_1/2_ = ln 0.5 /−*τ*. Finally, the diffusion coefficient D was measured using D = 0.88 * w^2^ / 4*τ*_1/2_ where w is the radius of the bleached area and τ the recovery time obtained by fitting the data.

## Acknowledgments

This work was supported by the DBT/Wellcome Trust India Alliance Fellowship [grant number: IA/I/15/1/501817] awarded to SR.

SR acknowledges and thanks Dr. Raj K. Ladher, NCBS, for generously donating plasmids of Cdh23 and Pcdh15 mammalian constructs. SR acknowledges the financial support provided by the DBT/Wellcome Trust India Alliance, Indian Institute of Science Education and Research Mohali (IISERM), and Centre for Protein Science Design and Engineering (CPSDE), Indian Institute of Science Education and Research Mohali. VK sincerely thank IISERM for financial support.

## Author Contributions

SR has supervised the project. VK did protein expression and purification performed the PICUP experiments in solution and on the membrane. VK, SKG, and TB performed FRAP on GUVs. VK and TB analysed FRAP data. VK and SR made all the figures. SR and VK wrote and edited the manuscript.

## Graphical Abstract

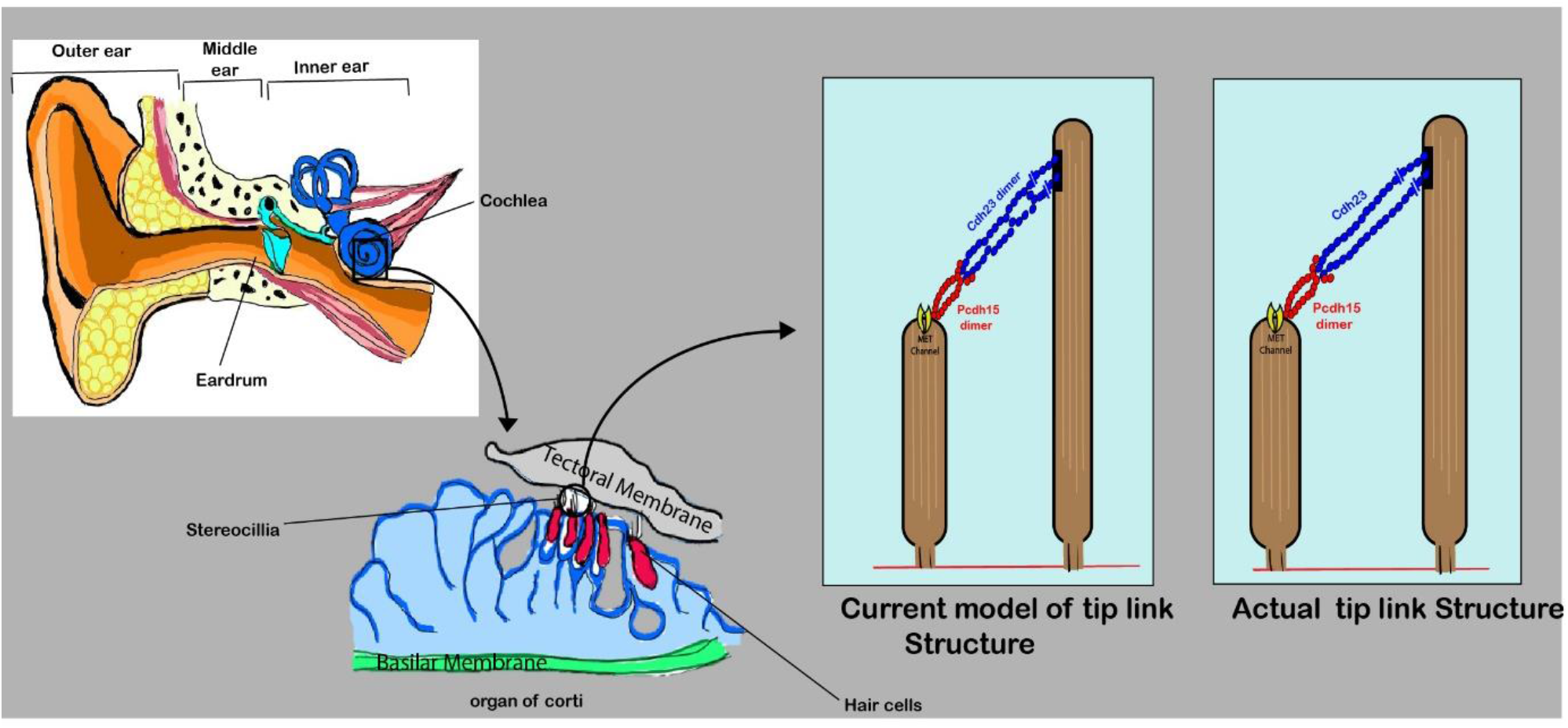

