## Supplementary file for "Redefining the structure of tip-links in hair-cells"

*Correspondence to Sabyasachi Rakshit

**This file includes:**

**Figure S1**

**Table S1**

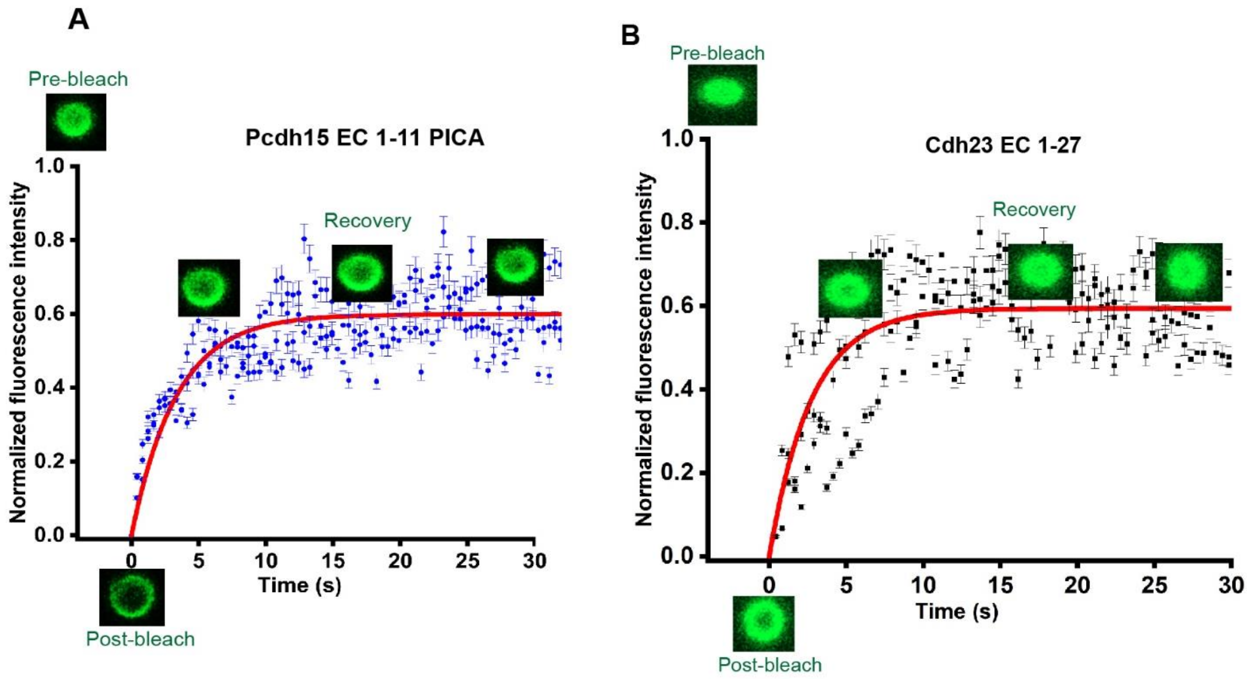


**Figure S1. Diffusion of proteins on GUVs tethered with Pcdh15 EC 1-11 PICA and Cdh23 EC 1-27:** The graph shows the recovery profile of normalized fluorescence intensity with time inside the photobleached region of (**A**) Pcdh15 EC 1-11 PICA tethered GUVs and **(B)** Cdh23 EC 1-27 anchored GUVs.Fluorescence images indicate sequences of FRAP events (i.e Pre – bleach, Post – bleach, and Recovery) on protein tethered GUVs. A solid red curve corresponds to the best fitting of fluorescence recovery over time The diffusion coefficient(D)was measured using $D = 0.88w^2 / 4\tau_{1/2}$ where w is the radius of the bleached part and the recovery time obtained by fitting the data (red curve). D measured for Pcdh15 EC 1-11 PICA and Cdh23 EC 1-27 is 0.361 ± 0.01 $\mu m^2$ /s and 0.23 ± 0.04 $\mu m^2$ /s respectively.

**Table S1. Diffusion coefficients measured for Pcdh15 EC 1-11 PICA and Cdh23 anchored to GUV membranes**

| **Protein anchored to GUV membrane** | **Diffusion coefficient ($\mu m^2$ /s)** |
|---|---|
| Pcdh15 EC 1-11 PICA | 0.361 ± 0.01 |
| Cdh23 EC1-27 | 0.23 ± 0.04 |
